# Complex Interplay between Serum and Fibroblasts in 3D Hepatocyte Co-culture

**DOI:** 10.1101/286088

**Authors:** Nikhil Mittal, Huan Li, Abhishek Ananthanarayanan, Hanry Yu

## Abstract

Previous studies have suggested that primary hepatocytes cultured *in vitro* undergo a rapid loss of function. On the other hand, in the clinic, drug induced liver injury typically manifests 5 days to 3 months after starting a medication. Thus, novel approaches that can maintain the function of primary human hepatocytes for longer durations of time may enable the development of improved *in vitro* assays for detecting hepatotoxicity. Previous studies have demonstrated that two-dimensional micro-patterning of hepatocytes with fibroblasts leads to improved maintenance of the hepatocyte phenotype relative to hepatocyte monocultures, in serum containing medium. Additionally, we, and others, have shown that three-dimensional culture of hepatocytes leads to enhanced function (in serum-free medium). In this study we wanted to (i) examine the effect of combining the above two approaches on hepatocyte function, and (ii) to further delineate the effect of serum on hepatocyte function. We developed a user-friendly and inexpensive approach for constructing layered spheroids. Similar to previous results in two-dimensional (2d) culture, we observed that 3d culture of hepatocytes alone (i.e. monoculture) in serum-containing medium led to an increase in the urea production rate, but near-complete loss of cytochrome activity in both lots of primary human hepatocytes (PHH) tested. In serum-free sandwich culture, cytochrome activity was maintained at the level observed in freshly thawed PHH for one lot, but almost completely lost in another lot. Spheroid culture of both lots of PHH in serum-free medium led to maintenance of CYP3A4 and CYP1A2 activity at the fresh thaw level, though CYP2B6 activity was reduced. In contrast to PHH monoculture, co-cultures of PHH with NIH 3T3 fibroblast cells benefitted from the presence of serum, and led to 3-5-fold increases in CYP activity relative to even serum-free spheroid monocultures. Layering of the fibroblasts did not result in improvements over mixed co-cultures. These results indicate the importance of appropriate serum-free monoculture control experiments in the evaluation of novel biomaterials and techniques for hepatocyte co-culture. Further, urea production and cytochrome production are decoupled; therefore, urea production is an insufficient readout when developing models for pharmaceutical applications.

## Introduction

### Motivation

Hepatotoxicity is one of the leading causes of the failure of drugs in clinical trials, and postmarket drug withdrawals. The ability to predict the potential hepatotoxicity of drug candidates earlier in development would result in enhanced patient safety. Additionally, this would lead to a reduction in the drug development cost, since late stage failures are associated with large monetary losses for pharmaceutical companies. Increasingly, these companies appear to be interested in performing in vitro toxicity screening during the early screening/discovery and lead optimization stages [1]. The ideal cell source for this application would be primary human hepatocytes (PHH) since they represent the cell type that is often affected as a result of drug induced liver injury (DILI). However, these cells typically undergo loss of function (sometimes referred to as de-differentiation) over a period of hours to days [2-5]. On the other hand, in the clinic, DILI typically manifests 5 days to 3 months after starting a medication [6]. Thus, novel approaches that can maintain the function of primary human hepatocytes for longer durations of time may enable the development of improved in vitro assays for detecting hepatotoxicity. Additionally, such systems would enable improved predictions of metabolism rates in slowly metabolized drugs.

Previous work has demonstrated that micro-patterned co-culture of PHH with 3T3 mouse fibroblast cells in serum-containing medium results in maintenance of the PHH phenotype over a 7-14 day period [7] (for further details see “Architecture” below, and especially [21]). However, pharmaceutical companies typically use serum-free medium formulations for hepatocyte studies such as toxicity testing [1] and testing the cytochrome induction potential of compounds. Therefore, we were additionally interested in understanding the interaction between serum and co-cultures.

### Previous work on the in vitro maintenance of hepatocyte function

#### Hepatocyte isolation

Prior to performing in vitro culture, hepatocytes must be isolated from the liver. Richert et al. [8] observed that the isolation process in rat livers led to ∼2-5 fold decreases in constitutive cytochrome (CYP)-dependent *activities*. Similarly, Saliem et al. determined that cryopreservation of PHH led to the decrease in the activity of several key cytochrome enzymes [9]. On the other hand, other studies found no significant change upon cryopreservation (reviewed in [10]). Also, a recent study (2015) using human livers found no significant loss in the level of most CYP proteins following hepatocyte isolation *and* cryopreservation [11]. However, the activity of these proteins was not evaluated in this study. Thus, for PHH it is currently somewhat unclear whether it is sufficient to maintain activity at the fresh thaw level, or whether it would be optimal to exceed the fresh thaw levels to recapitulate the levels in the parent liver.

#### Extracellular matrix configurations

Early work on hepatocyte de-differentiation typically used rat hepatocytes and focussed on the use of various extracellular matrix configurations (such as the collagen sandwich, collagen-matrigel sandwich etc.) for maintaining a differentiated state. Richert et al. [8] demonstrated that rat hepatocytes cultured on a single collagen layer remained attached until day 5 of culture, but ∼50% detached on day 6. However, Tuschl et al. [12] have demonstrated the ability to maintain such cultures for up to 10 days. Both groups demonstrated that hepatocytes cultured either in a sandwich configuration, or on a single layer on matrigel (Richert et al. only) remained attached until days 10–13 of culture. Importantly, Richert et al. observed that the isolation process led to ∼2-5 fold decreases in constitutive cytochrome (CYP)-dependent activities, and attachment led to a further ∼2-fold decrease in CYP activities. Following this stage, they, and Tuschl et al., demonstrated that CYP activities could be maintained by various matrix configurations, but they *did not recover to the values observed in freshly isolated cells*. On the other hand, UDP-glucuronosyl transferase and glutathione-dependent activities were equivalent in cultured hepatocytes and in rat livers. However, it is worth noting that there are large inter-individual differences in CYP expression across the human population [13, 14]. Thus, the ability to maintain CYP expression, albeit at a lower level, may still enable the in vitro modeling of human livers which have a lower level of CYP activity.

For primary human hepatocytes (PHH), Wilkening and Bader performed a time course analysis of gene expression in these hepatocytes, cultured within a collagen sandwich for 1 week [3]. They demonstrated that while certain CYPs (1A1, 1A2, 2C9, 2D6) could be maintained for a week, others (3A4, 2E1) declined. More recently, similar results were demonstrated for a collagen-matrigel sandwich by Bellwon et al. [4]. Khetani and Bhatia have also examined the expression of CYPs 2A6, 2B6, and 3A4 in PHH cultured in various matrix configurations and found that these CYPs could not be maintained, though it should be noted that their assays appear to have been performed in medium containing 10% fetal bovine serum (FBS) [7]. Studies have demonstrated that the presence of serum leads to reduction of CYP function in rat [12, 15] and human hepatocyte monocultures [16].

#### Co-culturing hepatocytes and other types of cells

Another approach to maintaining hepatocyte function in vitro has been to co-culture hepatocytes with other types of cells. The liver contains at least 5 cell types in addition to hepatocytes – sinusoidal endothelial cells, Kupffer cells, stellate cells, oval cells, and cholangiocytes. It is hypothesized that signals arising from these cells modulate the hepatocyte phenotype. Several earlier studies demonstrated that certain functions such as albumin secretion by PHH could be stabilized for several weeks in vitro by co-culturing PHH with various other cell types (reviewed in Bhatia et al. [17] and LeCluyse et al. [18]). Interestingly, as described in the above reviews, several groups found that even non-liver-associated cells such as murine embryonic fibroblasts (3T3) stabilized hepatocyte function. 3T3 cells are relatively easy to culture and appear to maintain their hepatocyte-supporting function for at least 12 passages. Thus, we chose to co-culture hepatocytes with these fibroblasts in our study, though we also examined co-cultures with adult human dermal fibroblasts (AHDF).

#### Architecture: Micropatterning and Dimensionality

In an interesting study published in 2008, Khetani et al. additionally demonstrated the importance of liver tissue architecture in the maintenance of certain hepatocyte functions [7]. The multiple cell types in the liver are not arranged randomly. Rather, the liver exhibits a complex architecture. In the mouse liver, ∼35% of the hepatocyte area in contact with other hepatocytes, and ∼50% in contact with sinusoidal endothelial cells (and the rest presumably in contact with other liver cell types) [19]. This study also demonstrated that this architecture is altered in liver carcinomas, with the percentages changing to ∼50% (hepatocyte-hepatocyte), and ∼40% (hepatocyte-sinusoid). Using in vitro micropatterning techniques, Khetani et al. demonstrated that optimization of the amount of heterotypic contact in vitro leads to optimal hepatocyte albumin and urea production. In terms of albumin production and urea secretion, the optimal geometry consisted of 500 µm wide circular “islands” of human hepatocytes i.e. approximately 250 hepatocytes per island (separated by 1200 µm (center-to-center spacing)), and surrounded by mouse fibroblast cells. Interestingly, this differed from the optimal island size for *rat* hepatocyte – mouse fibroblast co-cultures, which was only 36 µm, with only 1-4 cells per island [20]. 4 cells per island would indeed correspond with a ∼50% heterotypic interface. It will be interesting to know if the percent heterotypic interface differs between the human liver and the rat liver (i.e. in vivo). This may account for the difference in the optimal in vitro interface for human versus rat hepatocytes. However, it should be noted that a comparison of cytochrome activity in random versus patterned co-cultures was not performed.

Khetani and Bhatia also demonstrated that the function of PHH in their system was superior to that of PHH cultured in various matrix configurations. However, their assays appear to have been performed in medium containing 10% fetal bovine serum (FBS) [7]. Several studies (and our own results below) have demonstrated that the presence of serum leads to reduction of hepatocyte cytochrome function in rat [12, 15] and human hepatocyte monocultures [16]. Another more recent study from their lab is also very interesting, in that it shows that the activity of CYP enzymes in this system can vary with time in a highly non-linear manner (this study is performed in serum free medium) [21]. Finally, it is interesting that while contact appears to improve albumin production in hepatocyte-3T3 co-cultures [22], it inhibits albumin production in hepatocyte-endothelial cell co-cultures [23].

Another simple but important aspect of architecture is dimensionality. Organisms are three dimensional, but cell cultures are typically “two dimensional” i.e. they consist of monolayers of cells. In previous work, we [24] and others [18, 25] have found that the three dimensional culture of hepatocytes leads to improved maintenance of function. Other studies have also investigated 3d co-cultures [25-29]. However, in these studies the co-cultures were not organized/micro-patterned. Interestingly, a recent study that examined co-cultures of *iPS-derived* hepatocytes with adult human dermal fibroblasts found that organization led to *reduced* production of albumin [23].

#### Objectives for this study

In this work, we further explored the role of architecture, fibroblast type, and presence of serum on in vitro hepatocyte function, to rationally develop a 3D hepatocyte co-culture model. We developed a novel method to co-culture hepatocytes and fibroblasts as spheroids in which the relative distribution is either random (“mixed spheroids”), or non-random/ micropatterned. For the micropatterned configuration we layered fibroblasts around hepatocyte spheroids. We describe the results below.

## Results

### Rat spheroid formation in microwell plates

For describing the performance of layered co-culture spheroids (LCS) we decided to explore the use of a commercially available microwell plate (Elplasia Inc.). These 96-well plates contain 110 concave microwells per well. According to the manufacturer’s specifications, the microwell diameter at the surface is 500 µm, and the depth is 400 µm. The surface is coated with polyhydroxyethylmethacrylate (p-HEMA) to prevent cell adhesion.

Previous studies have demonstrated that rat hepatocyte spheroids with a diameter of 100 µm demonstrate optimal albumin production [30]. Therefore, we initially tried to obtain spheroids with a diameter of either 100 or 200 µm. Rat hepatocytes plated at 25, 000 cells/well assembled into a single spheroid per microwell over the course of 3 days (Figure 1a). The diameter of the spheroids was measured to be 119 ± 15 µm (Figure 1b and c). Based on this observation, we predicted that 200 µm spheres would require ∼25000 × 8 = 200, 000 cells/ well. However, we found that at this density the heptaocytes did not assemble into spheres even after 7 days in culture (Figure S1). Thus, for further studies with rat hepatocytes we used 25, 000 cells/well i.e. ∼250 hepatocytes per spheroid.

**Figure 1.**
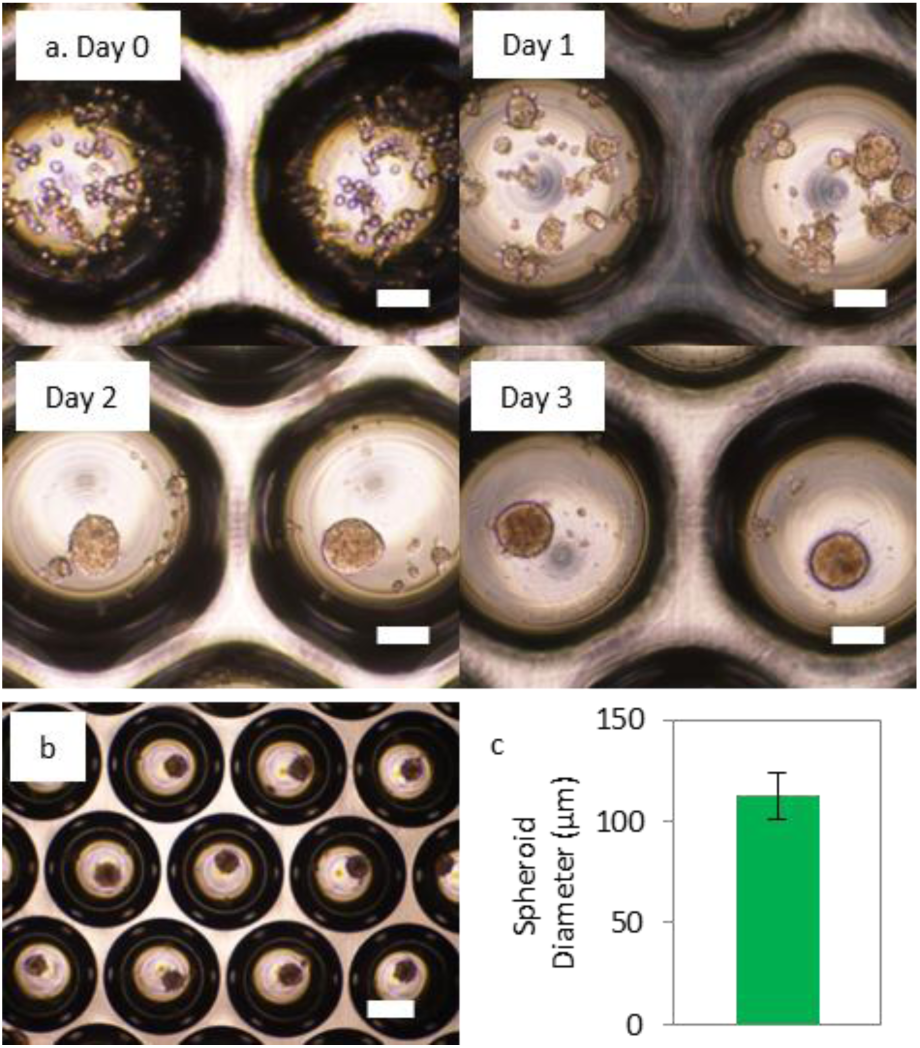
Formation of rat hepatocyte spheroids. a. Representative images of rat hepatocytes plated on a concave microwell array, Day 0 – Day 3 (∼250 hepatocytes per microwell). b. Representative lower magnification image of rat hepatocyte spheroids in a microwell array (day 3) c. Spheroid diameter (n=12 spheroids). Scale bars represent 100 µm (a) and 250 µm (b).

### Formation of mixed and layered co-culture spheroids (rat hepatocytes)

Next, we wanted to examine the function of three dimensional patterned co-cultures (Figure 2a). A previous study measured the surface area of NIH 3T3 fibroblasts to be ∼500 µm^2^ on soft substrates (3 kPa bulk stiffness) [31]. Therefore ∼80 fibroblasts are required to cover a sphere with a diameter of 119 µm (surface area ∼40,000 µm^2^). Since our system has 110 spheroids per well, this corresponds to 8800 fibroblast cells/well, or a hepatocyte: fibroblast of 25000:8800 i.e. approximately 3:1. Based on this estimate we explored ratios of 1:1, 3:2, 3:1, 6:1, and 10:1 (hepatocytes:fibroblasts).

**Figure 2.**
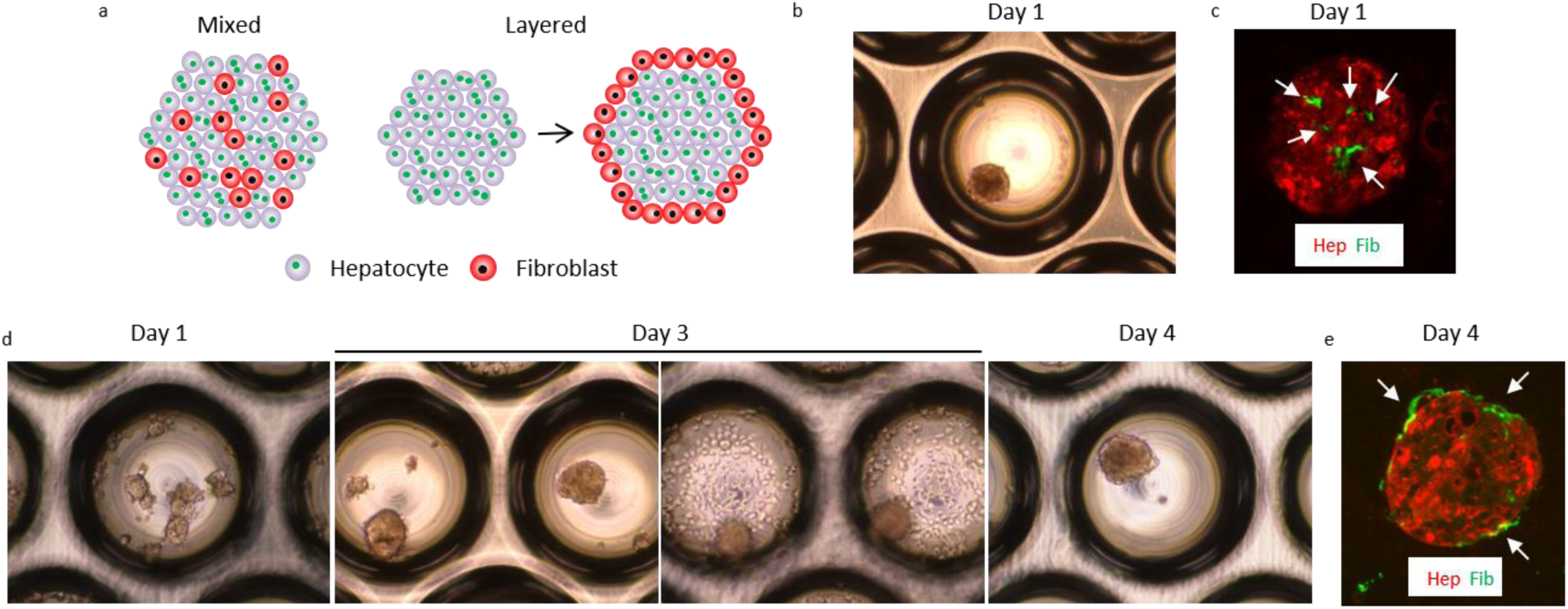
Formation of co-culture spheroids. a. Schematic demonstrating the two different co-culture configurations explored in this study. b. Representative phase contrast image of a mixed spheroid (6:1 hepatocytes: fibroblasts, Day 1). c. Section of a mixed spheroid (Day 4) stained for vimentin (green) and albumin (red). Arrows point to areas of vimentin staining (6:1 hepatocytes: fibroblasts). d. Formation of layered spheroids. Representative phase contrast images of: rat hepatocytes alone (Day 1, first panel), rat hepatocyte spheroids before (2^nd^ panel) and after (3^rd^ panel) the addition of adult human dermal fibroblast cells (Day 3, 1:1 hepatocytes: fibroblasts), co-culture spheroid (Day 4, 4^th^ panel). e. Section of a layered spheroid (Day 4) stained for vimentin (green) and albumin (red). Arrows point to areas of vimentin staining (6:1 hepatocytes: fibroblasts). Hep – hepatocyte, Fib – fibroblast.

All our experiments with rat hepatocytes were done in serum free William’s E medium (with epidermal growth factor). Rat hepatocytes mixed with either NIH 3T3 mouse fibroblasts or adult human dermal fibroblasts and plated into multiwell plates rapidly assembled into spheroids in only one day (Figure 2b) regardless of the co-culture ratio. To determine the relative location of fibroblasts and hepatocytes we sectioned and stained such spheroids for the hepatocyte marker albumin and the fibroblast marker vimentin. We observe that the fibroblasts are indeed distributed across the interior of the spheroid on Day 1 (Figure 2c, Figure 3c).

**Figure 3.**
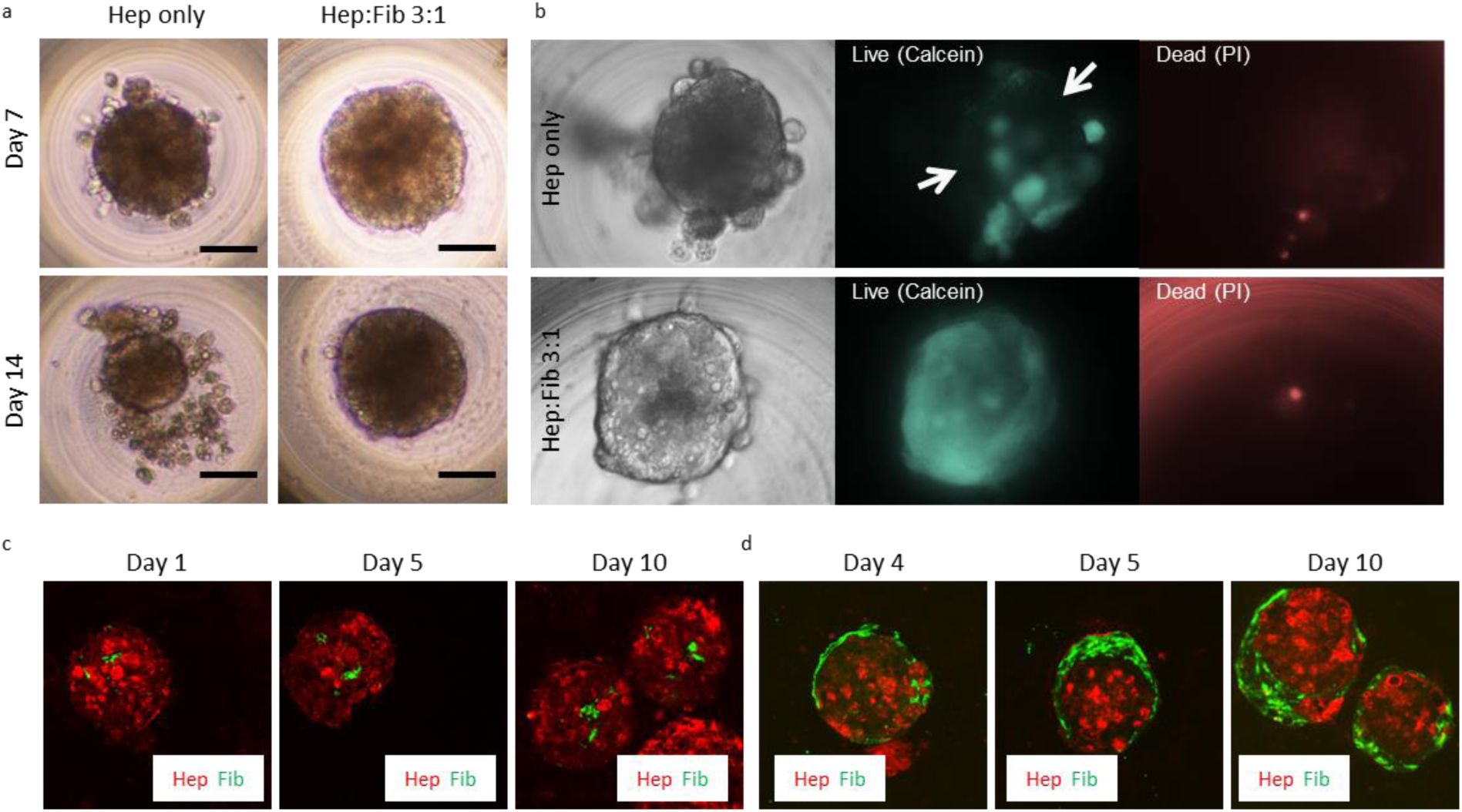
Maintenance and time-dependent organization of co-culture spheroids a. Representative images of mono and layered co-culture spheroids on days 7 & 14. Scale bars represent 50 µm. b. Representative images of mono and layered co-culture spheroids (day 14) stained with calcein and propidium iodide. The white arrows in the Live panel indicate regions with low fluorescence. c. Sections of mixed spheroids on days 1, 5, and 10 stained for vimentin (red) and albumin (green). d. Sections of layered spheroids on days 4 (first day after layering), 5, and 10 stained for vimentin (red) and albumin (green). Hep – hepatocyte, Fib – fibroblast.

For generating layered spheroids, we added fibroblasts to the culture following the formation of pure hepatocyte spheroids (Figure 2d – Day 3). Within a 24-hour period the added fibroblasts were incorporated into the spheroid (Supplementary Video 1, Figure 2d – Day 4). As expected, the fibroblasts adhered to the outer surface of the sphere (Figure 2e), although they were not always uniformly distributed (Figure 3d – Day 4 and Day 5).

A drawback of the current study design is that different populations of fibroblasts have to be used to generate the mixed versus layered spheroids because of the 3 days it takes the spheroids of pure hepatocytes to form. We tried centrifuging the plates as a way to enable more rapid formation of spheroids, but found no difference in the kinetics of spheroid formation (not shown). In the future, 3d printing may enable the simultaneous formation of mixed and layered spheroids, which would be a better controlled experiment.

### Maintenance and time-dependent organization of co-culture spheroids (rat hepatocytes)

Monoculture spheroids showed signs of disintegration by Day 14, while co-culture spheroids maintained their integrity (Figure 3a). We did not observe a higher number of dead cells in/around the monoculture spheroids (Figure 3b). However, they demonstrated very low esterase activity (Figure 3b – Live (Calcein), white arrows) suggesting that these cells were unhealthy. In mixed spheroids the fibroblasts remained distributed through the volume over the 10 day culture period (Figure 3c). In layered spheroids, the fibroblasts were largely maintained in a layer around the hepatocytes (Figure 3d).

### Functional characteristics of co-culture spheroids (rat hepatocytes)

We next measured urea and albumin production, and the expression of cytochrome enzymes for various co-culture ratios. At the high ratios of 1:1 or 3:2 (Hep:Fib), the spheroids aggregated into clumps via an unknown mechanism (Figure S2 a, b). These ratios were also associated with lower urea and albumin production (Figure S2 c, d), and so we did not examine these ratios in further assays.

For these assays, we primarily used adult human dermal fibroblasts (AHDF). For urea production on day 10, for layered spheroids, we observed an optimal ratio of 6:1 (Figure 4a). At this ratio urea production was enhanced relative to monoculture spheroids on day 2 or day 10 (Figure 4a). However, for mixed spheroids we observed no difference between monoculture and co-culture spheroids across all ratios (Figure 4b). Thus, spheroid architecture modulates urea production by rat hepatocytes (Figure 4c).

**Figure 4.**
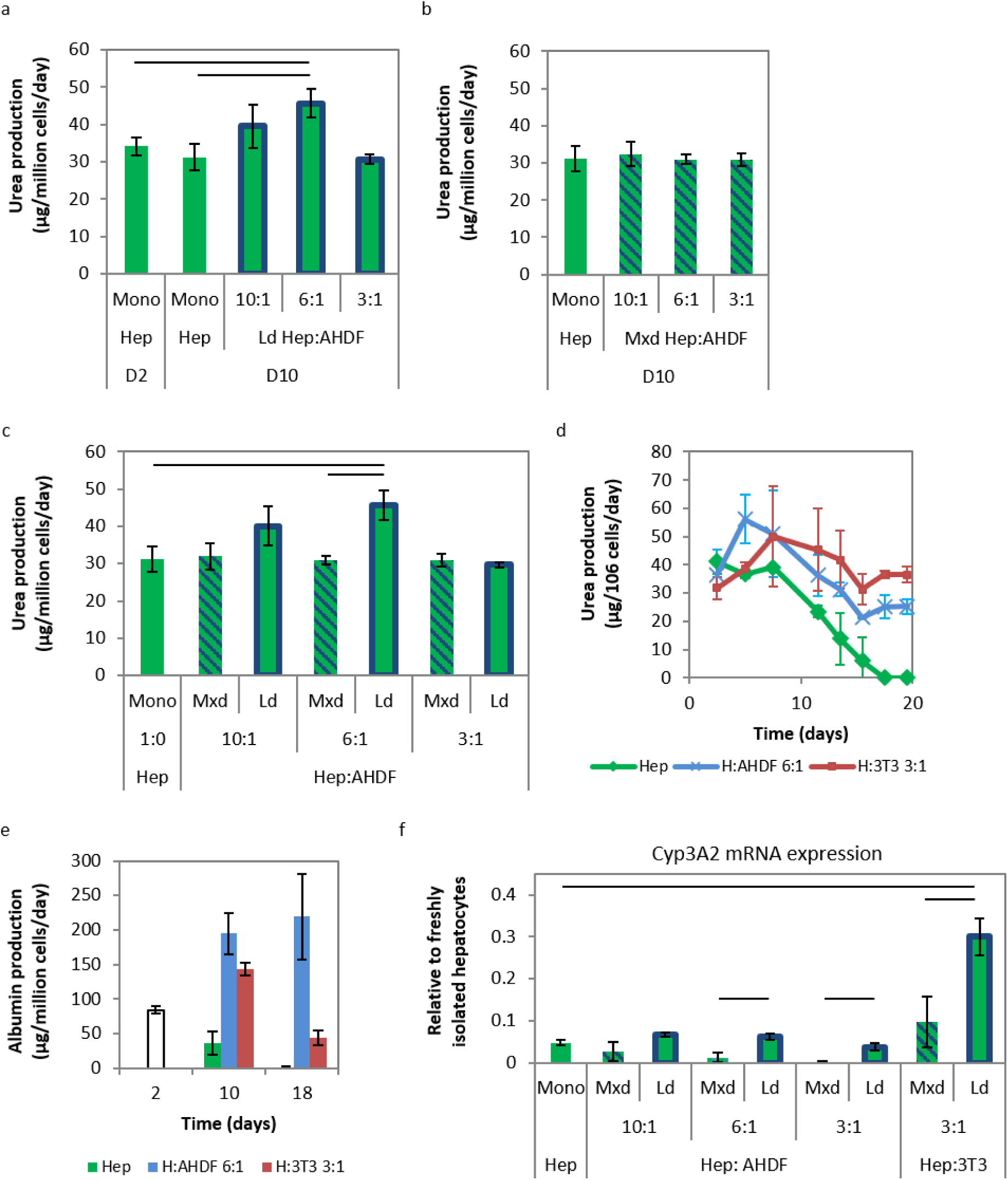
Functional characterization of rat hepatocyte spheroids a. Urea production in monoculture hepatocytes on day 2, monoculture and layered co-culture spheroids on day 10. b. Urea production by monoculture and mixed co-culture spheroids on day 10. c. Comparison of the urea production by mixed and layered co-culture spheroids on day 10. d. Time course of urea production by monoculture and layered co-culture spheroids. e. Time course of albumin production by monoculture and layered co-culture spheroids. f. Comparison of the Cyp3A2 mRNA expression in mixed and layered co-culture spheroids on day 10. AHDF – adult human dermal fibroblast, Hep – hepatocyte, Ld – layered, Mxd – mixed, 3T3 – NIH 3T3 fibroblast.

Next, we examined the time course of urea production in the 6:1 layered spheroid and monoculture spheroids (Figure 4d). We observed that monoculture spheroids maintained urea production for 8 days, after which there was a gradual decline in urea production, until it was lost at ∼ day 18. In contrast, urea production in layered 6:1 spheroids was non-monotonic and stabilized after day 16 at ∼25 µg/10^6^ cells/day, although the level was lower than the initial value (∼35 µg/10^6^ cells/day). A similar time course was obtained for layered spheroids made with 3T3 fibroblasts at a ratio of 3:1 (Hep:Fib), though the final value (35 µg/10^6^ cells/day) was higher than for AHDF.

We then examined albumin production in this system and found that there was no significant difference between mixed versus layered spheroids (not shown). However, while albumin production was lost in monoculture spheroids, it could be maintained in layered AHDF co-culture spheroids (again 6:1) (Figure 4e). Albumin production in 3T3 co-culture spheroids increased initially, but subsequently declined (Figure 4e).

Finally, we examined the expression of several cytochrome enzymes in this system. We observed significant variations in the expression of the housekeeping GAPDH (Glyceraldehyde 3-phosphate dehydrogenase, rat-specific primers). A similar observation has been made by Wilkening and Bader in the context of human hepatocytes [3]. On the other hand, as previously mentioned, we observed almost no cell death in our system (Figure 3b), therefore, the variation in the number of live cells per sample is less than 10 percent. Hence, we processed all the lysate for each sample in a similar manner and did not normalize to any housekeeping gene. In contrast to the production of urea, we observed that the presence of fibroblasts typically reduced the expression of cytochrome enzymes (mixed spheroids in Figure 4f and Figure S3). In some cases, layering resulted in the recovery of cytochrome expression, but expression was typically not higher relative to monoculture spheroids (see below). An exception to this trend was co-culture with 3T3 fibroblasts, which resulted in improved expression of Cyp3A2 and Cyp2B2 relative to monoculture spheroids (Figure 4f and Figure S3b).

It is worth noting that for rat hepatocytes plated even on collagen-coated polystyrene, a rapid loss of expression for several cytochrome enzymes occurs within 48 hours (Figure S3e), to ∼2-6% of the expression in freshly isolated cells. Similar observations have been made for PHH CYP activity [32]. For some of the CYPs (3A2, 2B2, 2D6) we observe a partial recovery in our system (Figure S3). However, the expression of other CYPs such as Cyp1A2 and Cyp2C11 reduced further over time (Figure S3).

### Formation of mixed and layered co-culture spheroids with human hepatocytes

A previous study demonstrated that the function of primary *human* hepatocytes (PHH) in micro-patterned co-cultured with 3T3 mouse fibroblasts was superior to that of PHH cultured in various matrix configurations [7]. However, it should be noted that their assays appear to have been performed in medium containing 10% fetal bovine serum (FBS). Studies have demonstrated that the presence of serum leads to reduction of hepatocyte function in rat [12, 15] and human hepatocyte (monocultures) [16]. Therefore, we investigated co-cultures of PHH with fibroblasts in the presence and absence of serum. Similar to rat hepatocytes, two lots of PHH studied formed spheroids within three days (Lot A – Figure 5a, Lot B – Figure S4a). Lot B showed lower viability relative to Lot A which resulted in considerable debris around the spheres (Figure S4a). This highlights one of the drawbacks of this system, which is that unlike 2d culture, the dead hepatocytes cannot be washed away since this would also lead to removal of the spheroids.

**Figure 5.**
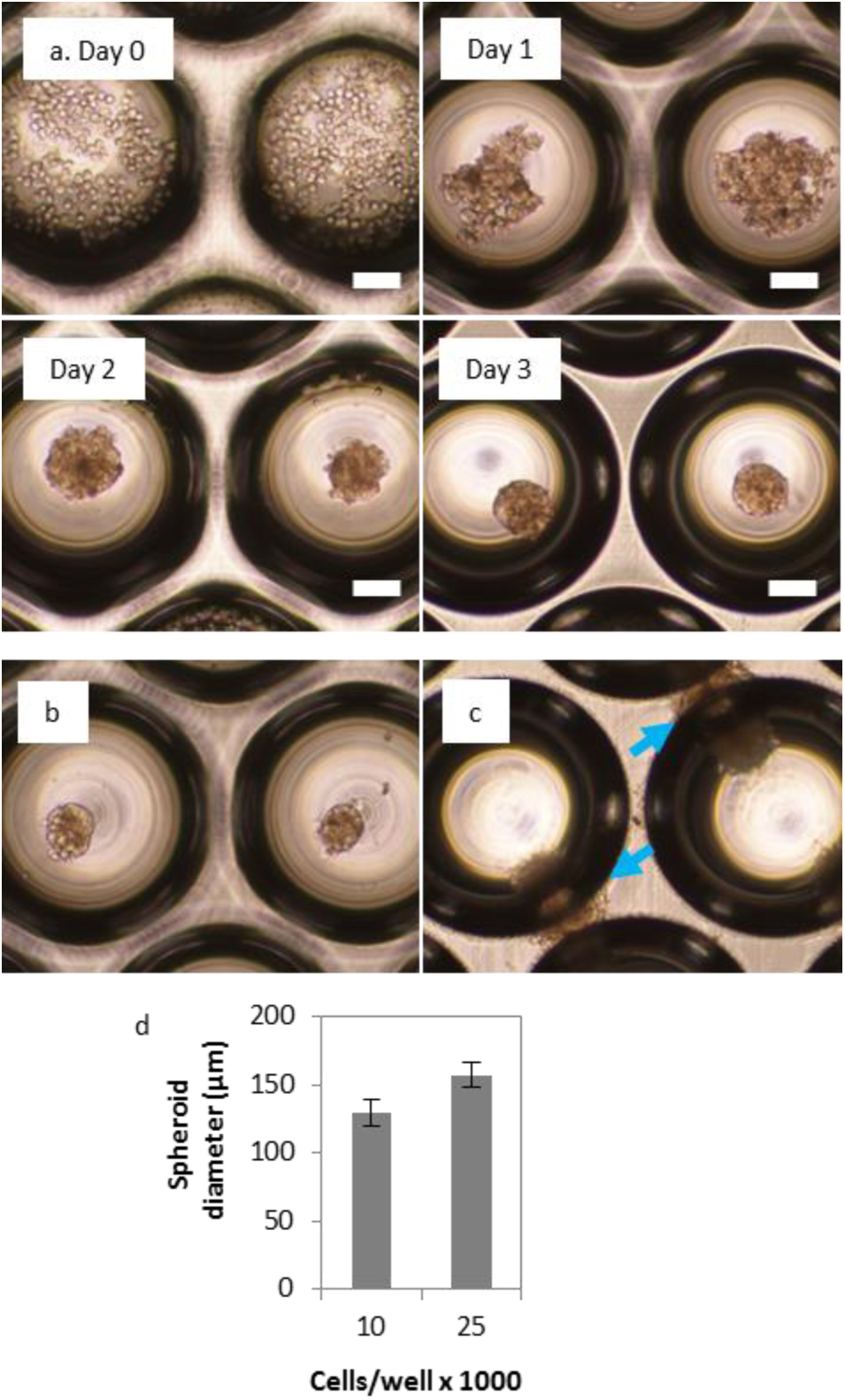
Formation of human hepatocyte spheroids. a. Representative images of human hepatocytes (Lot A) plated on a concave microwell array, Day 0 – Day 3, 250 hepatocytes per microwell (serum free medium). b & c. Spheroids on Day 7 for 10000 and 25000 cells per well respectively. The blue arrows point to spheroids that have migrated out of the microwells. d. Spheroid diameter for 10000 and 25000 cells per well (n = 4 spheroids). Scale bars represent 100 µm.

For 25000 hepatocytes per well, we observed that the spheroids migrated out of the microwells by day 7-10 (Figure 5c, Figure S4c). Hence, we reduced the cell number to 10,000 cells/well. At this density the spheroids were maintained within the microwells for the duration of the assay (14 days; Figure 5b, Figure S4b). The spheroid diameter for this density was 129.6 ± 9.7 µm.

Similar to co-cultures with rat hepatocytes, 3T3 and AHDF formed mixed and layered spheroids with PHH (Figure S5). Since CYP expression did not show a strong dependence on the co-culture ratio (Figure 4f, Figure S3) and due to the high cost of PHH, we only tested one ratio of PHH:fibroblasts. Since optimal urea production in co-cultures with rat hepatocytes was observed at a ratio of 6:1, we decided to use this ratio. However, in going from 25000 cells/spheroid to 10000 cells/spheroid, the volume of the spheroid was reduced by a factor of ∼ 25000/10000 = 2.5. But we wanted to preserve the amount of layering, and since the surface area ∼ radius^2^, the surface area would only have reduced by a factor of 2.5^2/3^ i.e. 1.84. Therefore, the new ratio used was (6/2.5) hepatocytes to (1/1.84) fibroblasts i.e. 4.4:1. To make the handling easier we used a ratio of 4:1. Additionally, since the most improvement in the level of Cyp3a2 was observed in co-cultures with 3T3 fibroblasts, we primarily focussed on this supporting cell type.

Again, similar to rat hepatocytes, by day 14, monoculture spheroids showed evidence of disintegration (Figure S6a, i), while co-culture spheroids maintained their structure (Figure S6b, j). Studies have demonstrated that the presence of serum leads to reduction of hepatocyte function in rat [12, 15] and human hepatocytes [16]. In agreement with these results, we observed that in the presence of serum, monoculture spheroids demonstrated a greater degree of disintegration (especially Lot B, Figure S6 k, d). On the other hand, co-culture spheroids maintained their structure, and were larger in (10%) serum-containing medium (Figure S6 e, i), likely due to the proliferation of fibroblasts [7]. Interestingly, for co-cultures in serum containing medium, by day 14 we observed a layer of fibroblasts covering the region between the wells (Figure S6f). We did not observe this layer in serum-free cultures (Figure S6c).

### Functional characteristics of co-culture spheroids (human hepatocytes)

Culture of PHH in serum containing medium led to enhanced urea production (Figure 6 a, e). However, consistent with previous reports that used hepatocyte monolayers [12, 15, 16], the presence of serum in the medium led to near-complete loss of CYP activity in monoculture spheroids (Figure 6). On the other hand, spheroid culture of both lots of PHH in *serum-free* medium resulted in maintenance of CYP3A4 and CYP1A2 activity at the fresh thaw level, though CYP2B6 activity did decrease. Thus, for serum monoculture, urea production is high, but cytochrome activity is very low. This indicates the need to use cytochrome activity as a readout when optimizing hepatocyte-based systems for pharmaceutical applications.

**Figure 6.**
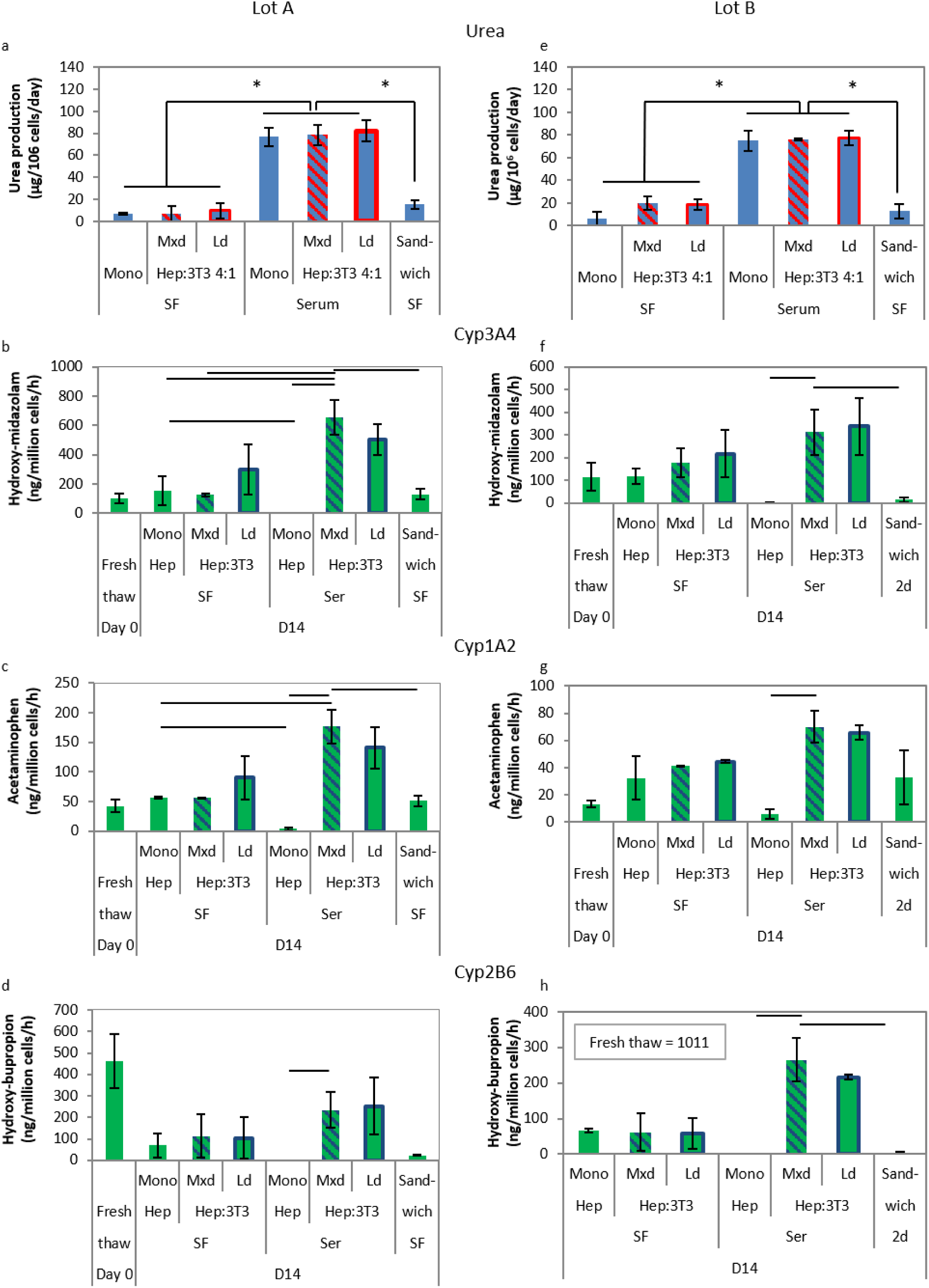
Function of mono- and co-culture spheroids for Lot A and B PHH on day 14. a and e. Urea production by mono- and co-culture spheroids on day 14 for Lot A (a) and Lot B (e). b, c, d. Cyp3A4, Cyp1A2, and Cyp2B6 activity for Lot A mono- and co-culture spheroids on day 14. Controls shown are the activity in freshly thawed cells (suspension) as well as hepatocytes in a collagen-matrigel sandwich. f, g, h. Cyp3A4, Cyp1A2, and Cyp2B6 activity for Lot B mono- and co-culture spheroids on day 14, along with suspension and sandwich values. Solid lines and asterisk indicate p<0.05. SF-serum-free medium, Ser – 10% serum-containing medium; Mxd – mixed co-culture; Ld – layered co-culture.

Next, we benchmarked PHH in our system to *serum free* PHH monoculture in 2 dimensions within a collagen-matrigel sandwich, a “standard” method used in the pharmaceutical industry (M. McMillian, personal communication). In serum-free sandwich culture, cytochrome activity was maintained at the level observed in freshly thawed PHH over 14 days for one lot (Lot A – Figure 6). For Lot B, activity could be maintained for 7 days (Figure S7), but was almost completely lost by D14 (Figure 6).

Interestingly, in contrast to the situation with PHH monoculture, co-cultures of PHH with NIH 3T3 fibroblast cells benefitted from the presence of serum, and 3-5-fold increases in CYP activity relative to (serum-free) spheroid monocultures were observed. But layering of the fibroblasts did not result in improvements over mixed co-cultures. For Lot A, pANOVA was 0.02, 0.01 and 0.07 for CYP3A4, CYP1A2, and CYP2B6 respectively. For Lot B, pANOVA was 0.09, 0.02 and 0.01 for CYP3A4, CYP1A2, and CYP2B6 respectively.

## Discussion

Primary hepatocytes cultured in vitro undergo loss of function over a period of hours to days. On the other hand, in the clinic, drug induced liver injury (DILI) typically manifests 5 days to 3 months after starting a medication. Thus, novel approaches that can maintain the function of primary human hepatocytes for longer durations of time may enable the development of improved in vitro assays for detecting hepatotoxicity. It is worth noting, however, that it is increasingly recognized that DILI typically falls into two categories – intrinsic or idiosyncratic [33]. Intrinsic DILI typically exhibits a dose-response relationship, and can often be detected via animal testing. However, in vitro testing may help reduce the associated cost. The potentially bigger challenge is to detect idiosyncratic DILI which can be caused by multiple mechanisms, including immune reactions to drug-protein complexes. Further, genotype could influence the development of toxicity. Thus, in effect, extension of the functional lifetime of hepatocytes in vitro represents one of a significant number of challenges that need to be solved to more accurately model clinical hepatotoxicity.

Previous studies have demonstrated that two-dimensional micro-patterning of hepatocytes with fibroblasts leads to improved maintenance of the hepatocyte phenotype relative to hepatocyte monocultures, in serum containing medium. Additionally, we, and others, have shown that three-dimensional culture of hepatocytes leads to enhanced function (in serum-free medium). In this study we examined the effect of combining the above two approaches on hepatocyte function, and further delineated the effect of serum on hepatocyte function. To enable the construction of layered spheroids we used a commercially available microwell plate. When compared to other spheroid platforms such as hanging drop plates, this platform offers the advantage of multiple spheroids per well. This enables the culture of 10-25000 hepatocytes/well, which is similar to the density typically used for hepatocyte culture (30,000 cells/well for rat hepatocytes and 50, 000 cells/well for human hepatocytes [34]. For studies of hepatotoxicity, a suitable number of cells/well is required for the development of appropriate concentrations of potentially toxic metabolites.

In future studies it would be interesting to further optimize the cell number per spheroid. In the current plate design, we observed that larger spheroids migrated out of the wells (Figure 5 and Figure S4). In future studies it may be possible to modify the design of the microwells to prevent spheroid migration, and this should enable the study of spheroids with larger numbers of cells. Indeed, a recent study used an approach similar to ours, to develop spheroids with 1500 PHH/spheroid [25]. Interestingly, this study demonstrated that 3d-cultured PHH closely resemble the in vivo liver at the proteome level relative to 2d cultures (no ECM overlay). Additionally, they demonstrate that the inter-sphere variability can be high ([25] – see for example Figure 4B). This highlights another advantage of a microwell system with multiple spheroids per well – due to averaging over multiple spheroids, interwell variability is reduced.

Rat and human primary hepatocytes formed spheroids in this system within 3 days (Figures 1 and 5). Upon addition of fibroblasts they were rapidly incorporated onto the surface of the spheroid (Figures 2 and S5). Layering of the fibroblasts led to an improvement in urea production. This is consistent with a previous 2d study that found that urea production increased more rapidly over time in patterned co-cultures relative to unpatterned ones [35]. Additionally, co-culture with 3T3 fibroblasts resulted in improved expression of certain CYPs (Cyp3A2 and Cyp2B2) relative to monoculture spheroids (Figure 4f and Figure S3b).

A drawback of the current study design is that different populations of fibroblasts have to be used to generate the mixed versus layered spheroids because of the 3 days it takes the spheroids of pure hepatocytes to form. We tried centrifuging the plates as a way to enable more rapid formation of spheroids, but found no difference in the kinetics of spheroid formation (not shown). In the future, 3d printing may enable the simultaneous formation of mixed and layered spheroids, which would be a better controlled experiment.

In our experiments with PHH, for one lot, we did observe that layering improved urea production at day 7 (Lot A – figure S7) relative to mixed co-culture. However, cytochrome activity was not significantly different in layered spheroids. Similarly, the previous 2d study demonstrated that micropatterning led to increased cumulative albumin and urea production relative to mixed co-cultures [7]. In future studies, it would be interesting to also assess cytochrome activity in 2d.

For PHH, we observed that co-culture results in 3-5-fold increases over monoculture, and over the fresh thaw levels. Since the relationship between the fresh thaw level of CYP activity and the activity levels in the donor liver is unknown for human PHH (see Introduction – hepatocyte isolation), it appears to be also unknown as to whether such an increase is truly desirable. However, the data from rat livers [8] suggests that a 2-5-fold decrease in activity occurs during hepatocyte isolation, and therefore a similar increase via co-culture would restore activity to the level in the native liver. For certain applications, however, it may be advantageous to accept a 3-5-fold lower activity level to avoid the additional complexity associated with culturing additional support cell types. Further, the fibroblasts proliferate and over time periods greater than the 2-week period used in this study, they may limit access of the hepatocytes to added drugs. Additionally, the central regions of the spheroids may become necrotic. Co-cultures with less proliferative non-parenchymal cells may mitigate this problem [25].

## Materials and Methods

### Cell culture

#### Rat hepatocytes

were isolated from male Wistar rats weighing 250-300 g using a modified in situ collagenase perfusion method [36]. Animals were handled according to the IACUC protocols approved by the IACUC committee of National University of Singapore. Cells were maintained with Williams’ E medium supplemented with 1 mg/mL BSA, 0.5 µg/mL of insulin, 100 nM dexamethasone, 50 ng/mL linoleic acid, 100 units/mL penicillin, and 100 µg/mL streptomycin and 10 ng/mL of EGF. They were incubated with 5% CO2 at 37^°^C and 95% humidity.

#### Human hepatocytes

Lot A was obtained from BD Biosciences (Donor HM P402), while Lot B was obtained from Life Technologies (Cellzdirect Lot HU 4227). Human hepatocytes were thawed in thawing/plating medium from Thermofisher (Williams E medium, 5% fetal bovine serum, 1 µM dexamethasone, 1% PenStrep, human recombinant insulin (4 µg/mL final concentration), 2 mM Glutamax, 15 mM HEPES. Human hepatocytes were cultured in cell maintenance medium from ThermoFisher (Williams’ E, 100 nM dexamethasone, 0.5% PenStrep, ITS (human recombinant insulin (6.25 µg/mL final concentration), human transferrin (6.25 µg/mL), selenous acid (6.25 ng/mL), bovine serum albumin (1.25 mg/mL), linoleic acid (5.35 µg/mL)), 2 mM Glutamax, 15 mM HEPES.

For microwell plates, rat hepatocytes were seeded in 100 µl and another 100 µl of medium was added the following day. To prevent the spheroids from coming out of the microwells, all medium change operations were performed slowly (∼4 seconds per operation, typically using an 8-channel pipette). After day 1, we performed half medium changes every day. Human hepatocytes with or without fibroblasts were seeded in 100 µl plating medium. 100 µl of maintenance medium was added the following day. After day 1, we performed half medium changes every day using maintenance medium. For layering, on Day 3 we added 100 µl of cell suspension containing fibroblasts instead of 100 µl of medium. From Day 4 onwards, we performed half medium changes every day as with the other wells.

#### NIH 3T3 fibroblasts and adult human dermal fibroblasts

(Lonza) were maintained in DMEM with 10% fetal bovine serum. Co-cultures were performed in Williams’ E medium (rat) or human hepatocyte maintenance medium as described above.

#### Microwell plates

were purchased from Elplasia Inc. (Japan, 2018 catalog number RB 500 400 NA).

#### Sandwich culture

96 well tissue culture polystyrene plates were incubated with collagen (5 µg/cm^2^ in 0.02N acetic acid) for one hour. PHH were plated as described above. The following day, the cells were overlaid with matrigel (Geltrex – Thermofischer) by replacing the plating medium with cold maintenance medium with 0.25 mg/ml matrigel. <u>On days 4, 8, and 12, the overlay was repeated.</u>

### Spheroid size measurements

Image analysis was performed using ImageJ – made binary, inverted, holes filled, then analysed using the “Analyse Particles” function.

### Imaging

Phase contrast imaging was performed using an Olympus CKX41 microscope and a Nikon Digital Sight DS-5M camera. Fluorescence images were acquired using an Olympus IV81 microscope and a Photometrics Cool Snap HQ2 camera.

### Sectioning and staining

Spheroids were fixed overnight in 4% paraformaldehyde (∼100 spheroids per Eppendorf tube with 0.5 ml paraformaldehyde). The following day they were washed 3 times for >5 minutes each with 1.2 ml PBS/tube. Embedding and sectioning were performed by the Advanced Molecular Pathology Laboratory at the Institute of Molecular and Cellular Biology (Singapore). Prior to embedding into paraffin, the spheroids were re-suspended in 4% low melting point agarose in phosphate buffered saline and the resulting mixture was allowed to set. After rehydrating the slides (2X xylene, 2X ethanol, 1X 95% ethanol in DI water, 1X 70% ethanol, 1X 50% ethanol, PBS), the sections were blocked using 1% triton, 2% BSA in PBS. Primary staining was performed overnight at 4^°^C. Antibodies used were monoclonal rabbit anti-vimentin (AbCam ab92547, 1:200) and sheep anti-rat-albumin (Bethyl Laboratories, 1:200). Secondary antibodies were used at a concentration of 1:400.

### Albumin and Urea measurements

Albumin concentrations were measured using albumin enzyme-linked immunosorbent assay (ELISA) quantitation kit (Bethyl Laboratories, Inc., Montgomery, TX, USA) as per the manufacturer’s protocol.

Urea production was measured using a commercially available kit (Urea Nitrogen Direct kit, Standbio Laboratory, Boerne, TX, USA). Briefly, 20 µl of supernatant from cell culture media was heated with a 150 µl 2:1 mixture of acid reagent: color reagent at 90°C for 30 minutes. The reaction was stopped by cooling on ice and the optical density was measured at 520nm after allowing the mixture to stand at room temperature for at least 20 minutes.

The urea/albumin produced in a 24 hour period was calculated by measuring the urea and albumin concentration in medium collected 24 hours apart. Then, since half medium changes were performed, the amount produced was calculated as: 190 µl × DayN concentration – 100 µl × DayN-1 concentration. We used 190 µl since we observed 10 µl evaporation per day (per well).

### Gene expression analysis

RNA was extracted from cells using a commercially available kit (RNeasy Plus Mini kit, Qiagen). Conversion to cDNA was performed using iScript (Bio-Rad). qPCR was performed on an Applied Biosystems 7500 Fast real time qPCR system using the FastStart Universal SYBR Green Master Mix (Roche).

### Cytochrome activity

PHH were incubated with a mixture of 100 µl of 5µM midazolam and 200 µM phenacetin for 2 hours. After collecting the supernatant we added 10 ng of acetaminophen-D4 as an internal standard and dried the samples using a concentrator (Eppendorf 5301, Hamburg, Germany) under vacuum. The dried residues were reconstituted using methanol containing 0.1% formic acid, vortexed, sonicated for 3 minutes, and centrifuged at 10^4^ rpm for 10 min. The supernatants were then used for measurement by LC/MS system (LC: 1100 series, Agilent, Singapore; MS: LCQ Deca XP Max, Finnigan, Singapore) with 100 _ 3.0 mm onyx-monolithic C18 column (Phenomenex, CA, USA). The mobile phase consisted of solvent A (0.1% formic acid in water) and solvent B (0.1% formic acid in methanol) with a flow rate of 0.8 ml/min. The elution scheme for the measurement of acetaminophen and 10-OHemidazolam involved solvent B which was gradually increased from 6% to 90% over 6 min. For OH-bupropion measurement, solvent B was gradually increased from 10 to 90% over 6 min. The MS parameter settings were as follows: spray voltage 5 kV; sheath gas flow rate: 80; auxiliary gas flow rate: 20; capillary temperature: 350 ^o^C; tube lens: 45 V; and capillary voltage: 30 V.

## Acknowledgements

This work is supported in part by the Institute of Bioengineering and Nanotechnology, Biomedical Research Council, A*STAR, Joint Council Office grant IBN/13-J51002, a NMRC grant R-185-000-294-511 and SMART BioSyM and Mechanobiology Institute of Singapore (R-714-001-003-271) funding to HYU. We thank Dr. Farah Tasnim and Dr. McMillian (Invitrocue Pte. Ltd.) for useful discussions. We would also like to thank Kapish Gupta of the Mechanobiology Institute (Singapore) for assistance with time lapse imaging and the Advanced Molecular Pathology Laboratory at the Institute of Molecular and Cellular Biology (Singapore) for assistance with sectioning the spheroids.

## References

1. Schadt, S., et al., Minimizing DILI risk in drug discovery – A screening tool for drug candidates. Toxicol In Vitro, 2015. 30(1 Pt B): p. 429–37.

2. Bonzo, J.A., et al. Long Term Hepatocyte Culture for Drug Safety Assessment Screening. 2014 [cited 2016 3/31/2016]; Available from: https://www.thermofisher.com/content/dam/LifeTech/global/life-DF-10-14/Long%20Term%20Hepatocyte%20Culture%20for%20Drug%20Safety%20Assessment%20Screening.pdf.

3. Wilkening, S. and A. Bader, Influence of culture time on the expression of drug-metabolizing enzymes in primary human hepatocytes and hepatoma cell line HepG2. J Biochem Mol Toxicol, 2003. 17(4): p. 207–13.

4. Bellwon, P., et al., Kinetics and dynamics of cyclosporine A in three hepatic cell culture systems. Toxicol In Vitro, 2015. 30(1 Pt A): p. 62–78.

5. Lauschke, V.M., et al., Massive rearrangements of cellular MicroRNA signatures are key drivers of hepatocyte dedifferentiation. Hepatology, 2016. 64(5): p. 1743–1756.

6. Uetrecht, J., Idiosyncratic drug reactions: current understanding. Annu Rev Pharmacol Toxicol, 2007. 47: p. 513–39.

7. Khetani, S.R. and S.N. Bhatia, Microscale culture of human liver cells for drug development. Nat Biotechnol, 2008. 26(1): p.120–6.

8. Richert, L., et al., Evaluation of the effect of culture configuration on morphology, survival time, antioxidant status and metabolic capacities of cultured rat hepatocytes. Toxicol In Vitro, 2002. 16(1): p. 89–99.

9. Saliem, M., et al., Improved cryopreservation of human hepatocytes using a new xeno free cryoprotectant solution. World J Hepatol, 2012. 4(5): p. 176–83.

10. Li, A.P., Human hepatocytes: isolation, cryopreservation and applications in drug development. Chem Biol Interact, 2007. 168(1): p. 16–29.

11. Vildhede, A., et al., Comparative Proteomic Analysis of Human Liver Tissue and Isolated Hepatocytes with a Focus on Proteins Determining Drug Exposure. J Proteome Res, 2015. 14(8): p. 3305–14.

12. Tuschl, G., et al., Serum-free collagen sandwich cultures of adult rat hepatocytes maintain liver-like properties long term: a valuable model for in vitro toxicity and drug-drug interaction studies. Chem Biol Interact, 2009. 181(1): p. 124–37.

13. Snawder, J.E. and J.C. Lipscomb, Interindividual variance of cytochrome P450 forms in human hepatic microsomes: correlation of individual forms with xenobiotic metabolism and implications in risk assessment. Regul Toxicol Pharmacol, 2000. 32(2): p. 200–9.

14. Amaral, K.B., et al. Enzyme Activity and Transporter Facilitated Uptake Comparisons: Multi-donor Pooled Cryopreserved Hepatocytes (HEP10™) to Single Donor Constituent Lots. 2012 [cited 2016 4/1/2016]; Available from: https://www.thermofisher.com/content/dam/LifeTech/migration/files/drug-discovery-development/pdfs.par.1689.file.dat/issx%20hep10%20poster%20final%2012oct12.pdf.

15. Hammond, A.H. and J.R. Fry, Effect of serum-free medium on cytochrome P450- dependent metabolism and toxicity in rat cultured hepatocytes. Biochem Pharmacol, 1992. 44(7): p. 1461–4.

16. Takeba, Y., et al., Comparative study of culture conditions for maintaining CYP3A4 and ATP-binding cassette transporters activity in primary cultured human hepatocytes. J Pharmacol Sci, 2011. 115(4): p. 516–24.

17. Bhatia, S.N., et al., Effect of cell-cell interactions in preservation of cellular phenotype: cocultivation of hepatocytes and nonparenchymal cells. FASEB J, 1999. 13(14): p. 1883–900.

18. LeCluyse, E.L., et al., Organotypic liver culture models: meeting current challenges in toxicity testing. Crit Rev Toxicol, 2012. 42(6): p. 501–48.

19. Hoehme, S., et al., Prediction and validation of cell alignment along microvessels as order principle to restore tissue architecture in liver regeneration. Proc Natl Acad Sci U S A, 2010. 107(23): p. 10371–6.

20. Bhatia, S.N., et al., Probing heterotypic cell interactions: hepatocyte function in microfabricated co-cultures. J Biomater Sci Polym Ed, 1998. 9(11): p. 1137–60.

21. Lin, C., et al., Prediction of Drug Clearance and Drug-Drug Interactions in Microscale Cultures of Human Hepatocytes. Drug Metab Dispos, 2016. 44(1): p. 127–36.

22. Hui, E.E. and S.N. Bhatia, Micromechanical control of cell-cell interactions. Proc Natl Acad Sci U S A, 2007. 104(14): p. 5722–6.

23. Stevens, K.R., et al., InVERT molding for scalable control of tissue microarchitecture. Nat Commun, 2013. 4: p. 1847.

24. Nugraha, B., et al., Galactosylated cellulosic sponge for multi-well drug safety testing. Biomaterials, 2011. 32(29): p. 6982–94.

25. Bell, C.C., et al., Characterization of primary human hepatocyte spheroids as a model system for drug-induced liver injury, liver function and disease. Sci Rep, 2016. 6: p. 25187.

26. Kostadinova, R., et al., A long-term three dimensional liver co-culture system for improved prediction of clinically relevant drug-induced hepatotoxicity. Toxicol Appl Pharmacol, 2013. 268(1): p. 1–16.

27. Lu, H.F., et al., Three-dimensional co-culture of rat hepatocyte spheroids and NIH/3T3 fibroblasts enhances hepatocyte functional maintenance. Acta Biomater, 2005. 1(4): p. 399–410.

28. Seo, S.J., et al., Enhanced liver functions of hepatocytes cocultured with NIH 3T3 in the alginate/galactosylated chitosan scaffold. Biomaterials, 2006. 27(8): p. 1487–95.

29. Vernetti, L.A., et al., A human liver microphysiology platform for investigating physiology, drug safety, and disease models. Exp Biol Med (Maywood), 2016. 241(1): p. 101–14.

30. Glicklis, R., J.C. Merchuk, and S. Cohen, Modeling mass transfer in hepatocyte spheroids via cell viability, spheroid size, and hepatocellular functions. Biotechnol Bioeng, 2004. 86(6): p. 672–80.

31. Rajagopalan, P., et al., Direct comparison of the spread area, contractility, and migration of balb/c 3T3 fibroblasts adhered to fibronectin- and RGD-modified substrata. Biophys J, 2004. 87(4): p. 2818–27.

32. Ulvestad, M., et al., Drug metabolizing enzyme and transporter protein profiles of hepatocytes derived from human embryonic and induced pluripotent stem cells. Biochem Pharmacol, 2013. 86(5): p. 691–702.

33. Uetrecht, J. and D.J. Naisbitt, Idiosyncratic adverse drug reactions: current concepts. Pharmacol Rev, 2013. 65(2): p. 779–808.

34. LeCluyse, E.L., Human hepatocyte culture systems for the in vitro evaluation of cytochrome P450 expression and regulation. Eur J Pharm Sci, 2001. 13(4): p. 343–68.

35. Bhatia, S.N., et al., Microfabrication of hepatocyte/fibroblast co-cultures: role of homotypic cell interactions. Biotechnol Prog, 1998. 14(3): p. 378–87.

36. Seglen, P.O., Preparation of isolated rat liver cells. Methods Cell Biol, 1976. 13: p. 29–83.

