## Supplementary Materials for "Complex Interplay between Serum and Fibroblasts in 3D Hepatocyte Co-culture"

**Supplementary Figures**

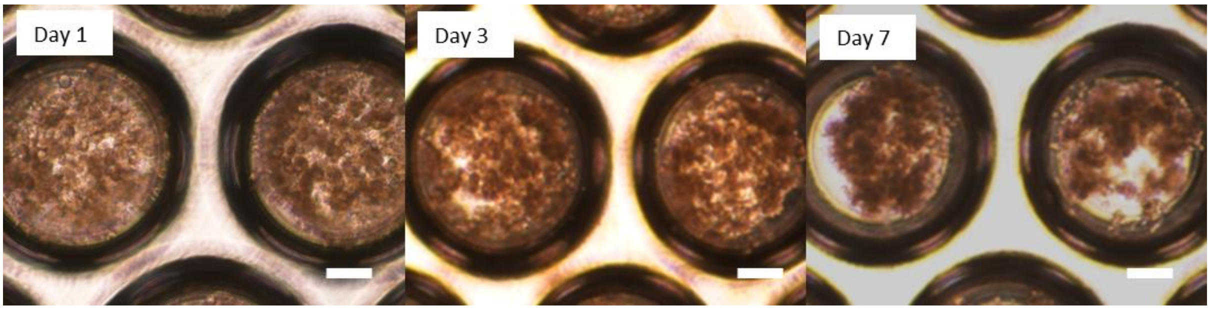


**Figure S1.** 2000 hepatocytes per microwell would theoretically form a spheroid with a diameter of 200 µm. However, at this density the hepatocytes did not condense into spheroids. The figure shows representative images of hepatocytes plated at this density on the microwell array (Day 0, 3, 7). Scale bars represent 100 µm.

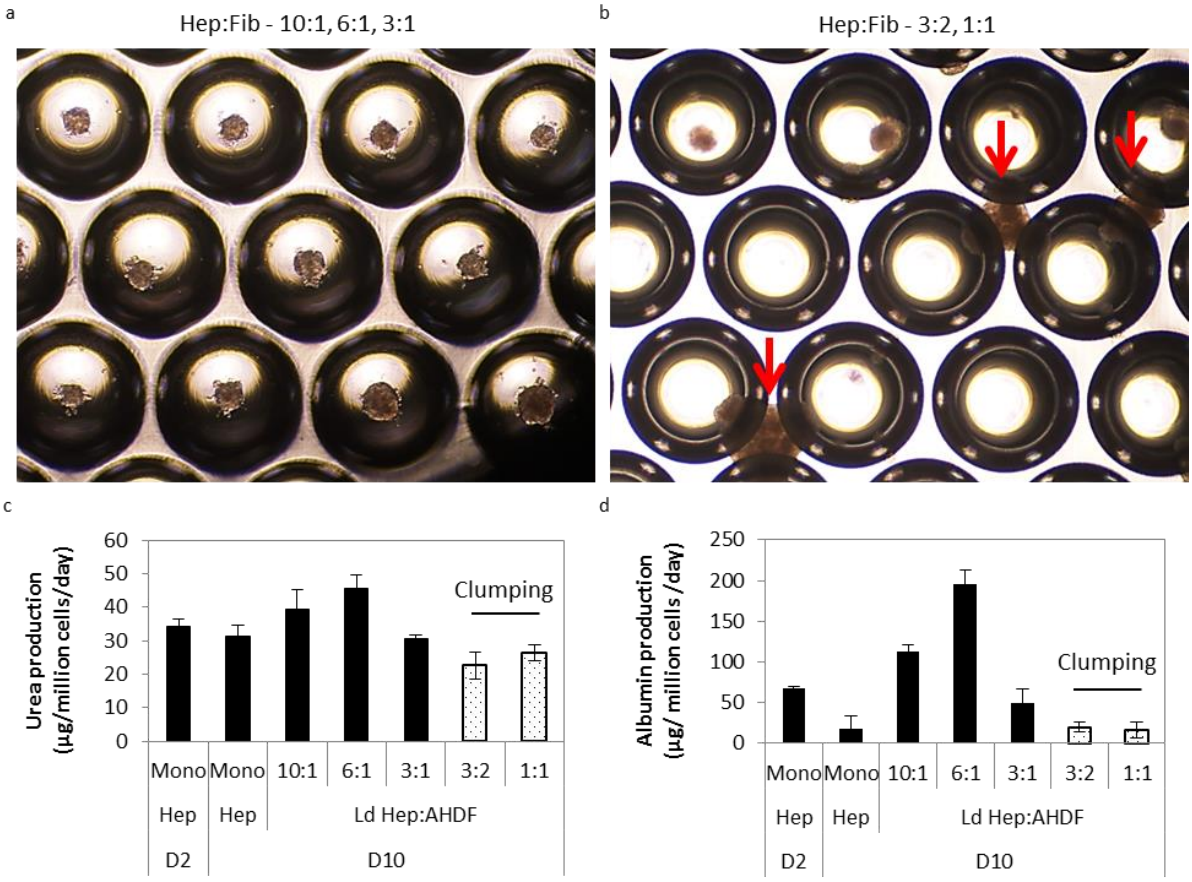


**Figure S2**. Optimization of the co-culture ratio. a. Co-culture ratios of 10:1 (hepatocytes: fibroblasts), 6:1 and 3:1 led to discrete layered spheroids within microwells. b. Increasing the fibroblast percentage i.e. co-culture ratios of 3:2 (hepatocytes: fibroblasts) and 1:1 led to the clumping of layered spheroids. The red arrows point to such clumps. c & d. In initial experiments we observed that this clumping led to lower production of urea (c) and albumin (d). Thus, only co-culture ratios of 10:1, 6:1, and 3:1 were investigated further. AHDF – adult human dermal fibroblast, Hep – hepatocyte, Ld – layered.

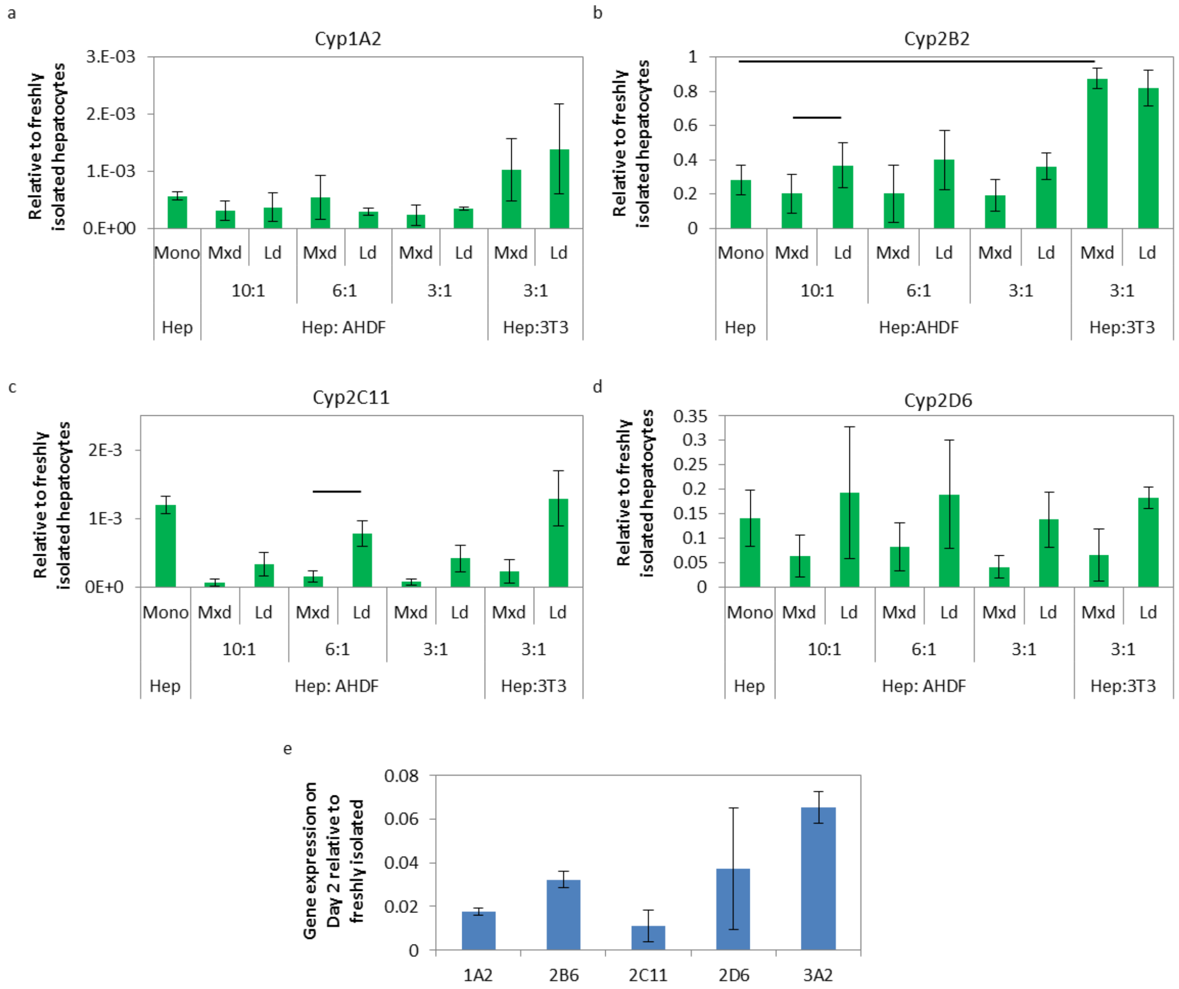


**Figure S3.** a-d. Comparison of the cytochrome mRNA expression in mixed and layered co-culture spheroids on day 10. AHDF – adult human dermal fibroblast, Hep – hepatocyte, Ld – layered, Mxd – mixed, 3T3 – NIH 3T3 fibroblast. e. Cytochrome mRNA expression in rat hepatocytes after 48 hours of culture on collagen-coated polystyrene, relative to freshly isolated cells.

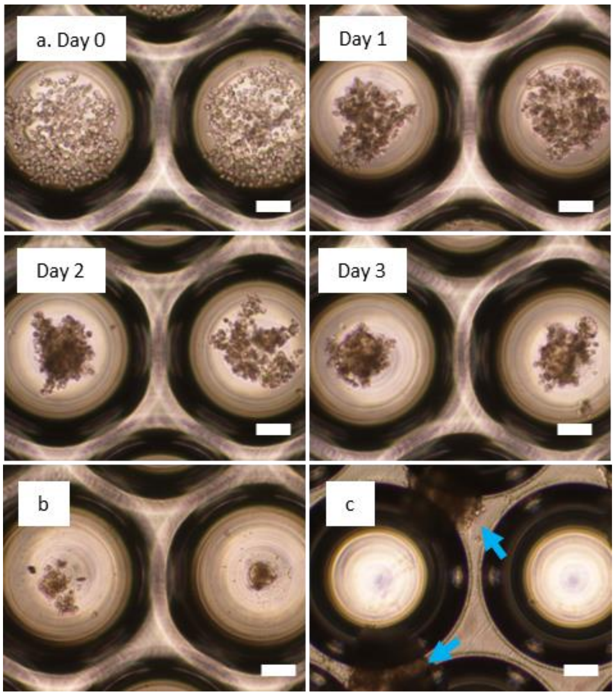


**Figure S4**. Formation of human hepatocyte spheroids (Lot B). a. Representative images of human hepatocytes plated on a concave microwell array, Day 0 – Day 3, 250 hepatocytes per microwell (serum free medium). b & c. Spheroids on Day 7 for 10000 and 25000 cells per well. Scale bars represent 100 μm.

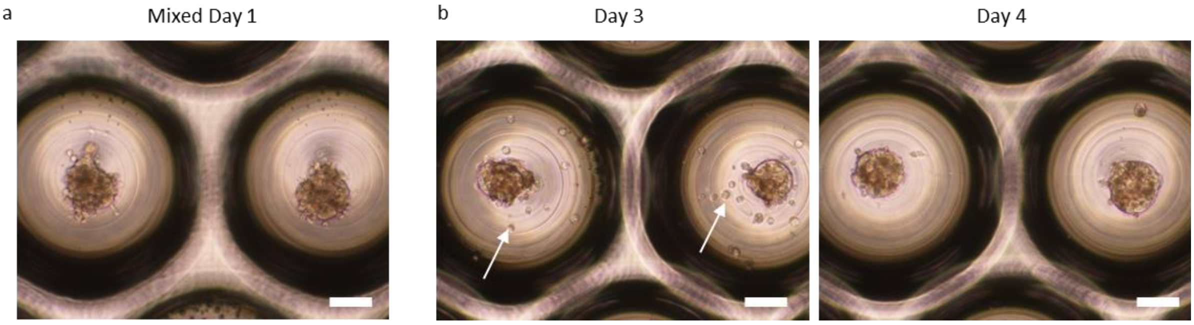


**Figure S5**. Formation of co-culture spheroids with human hepatocytes (Lot A). a. Representative phase contrast image of a mixed spheroid (4:1 hepatocytes: 3T3 fibroblasts; Day 1). b. Formation of layered spheroids. Representative phase contrast images of: human hepatocytes after the addition of fibroblast cells (Day 3, 4:1 hepatocytes: 3T3 fibroblasts), co-culture spheroid (Day 4, 2$^{nd}$ panel). The arrows point to fibroblast cells. Scale bar represents 100 μm.

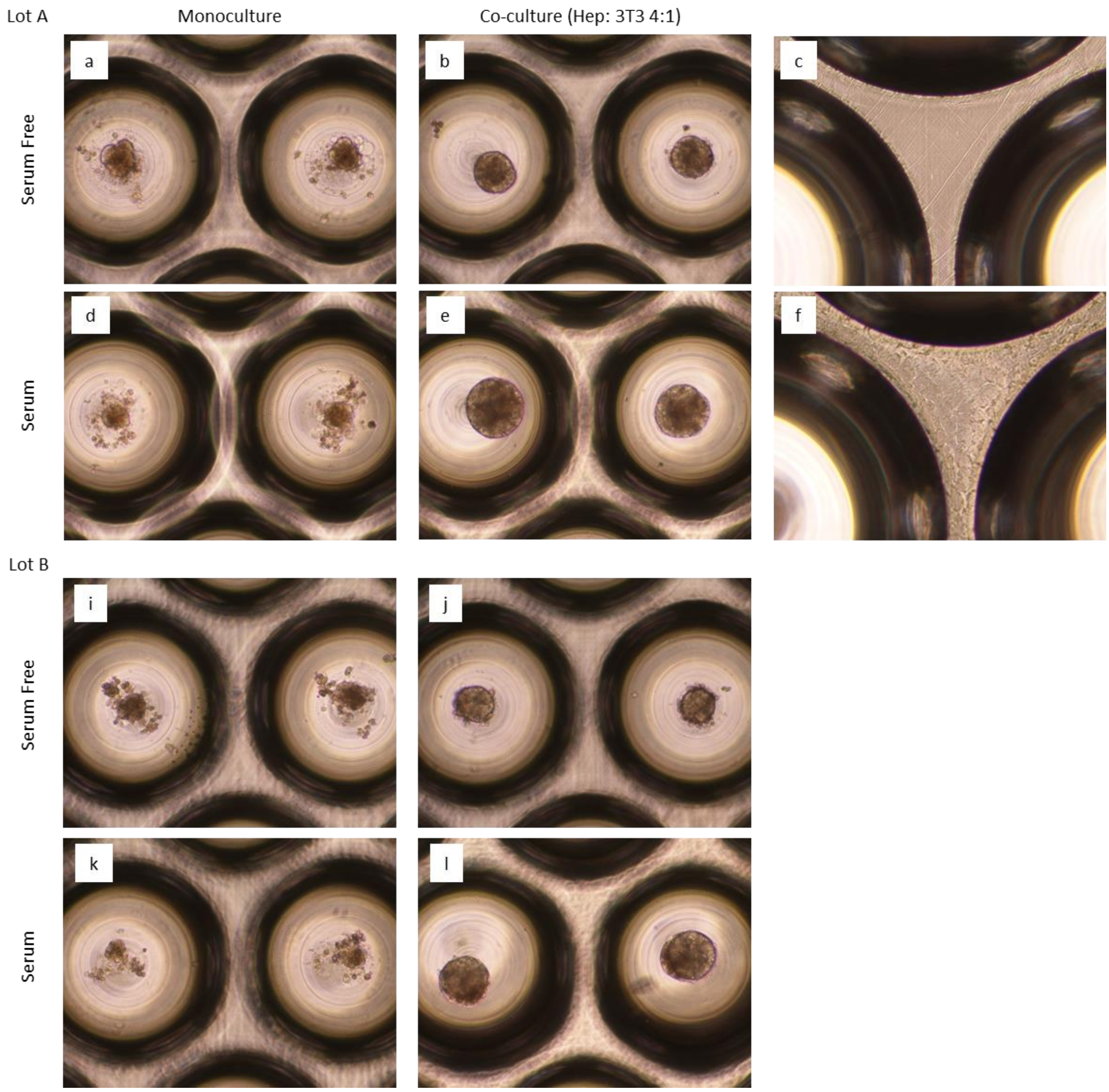


**Figure S6.** Maintenance of spheroids in our system. Phase contrast images of spheroids in microwell arrays. a-f Lot A human hepatocytes. i-l Lot B human hepatocytes. Monoculture spheroids in serum free (a, i) and (10%) serum containing medium (d, k). Co-culture (layered) spheroids in serum free (b, j) and (10%) serum containing medium (e, l). Inter-well region for co-cultures in serum free (c) and (10%) serum containing medium (f).

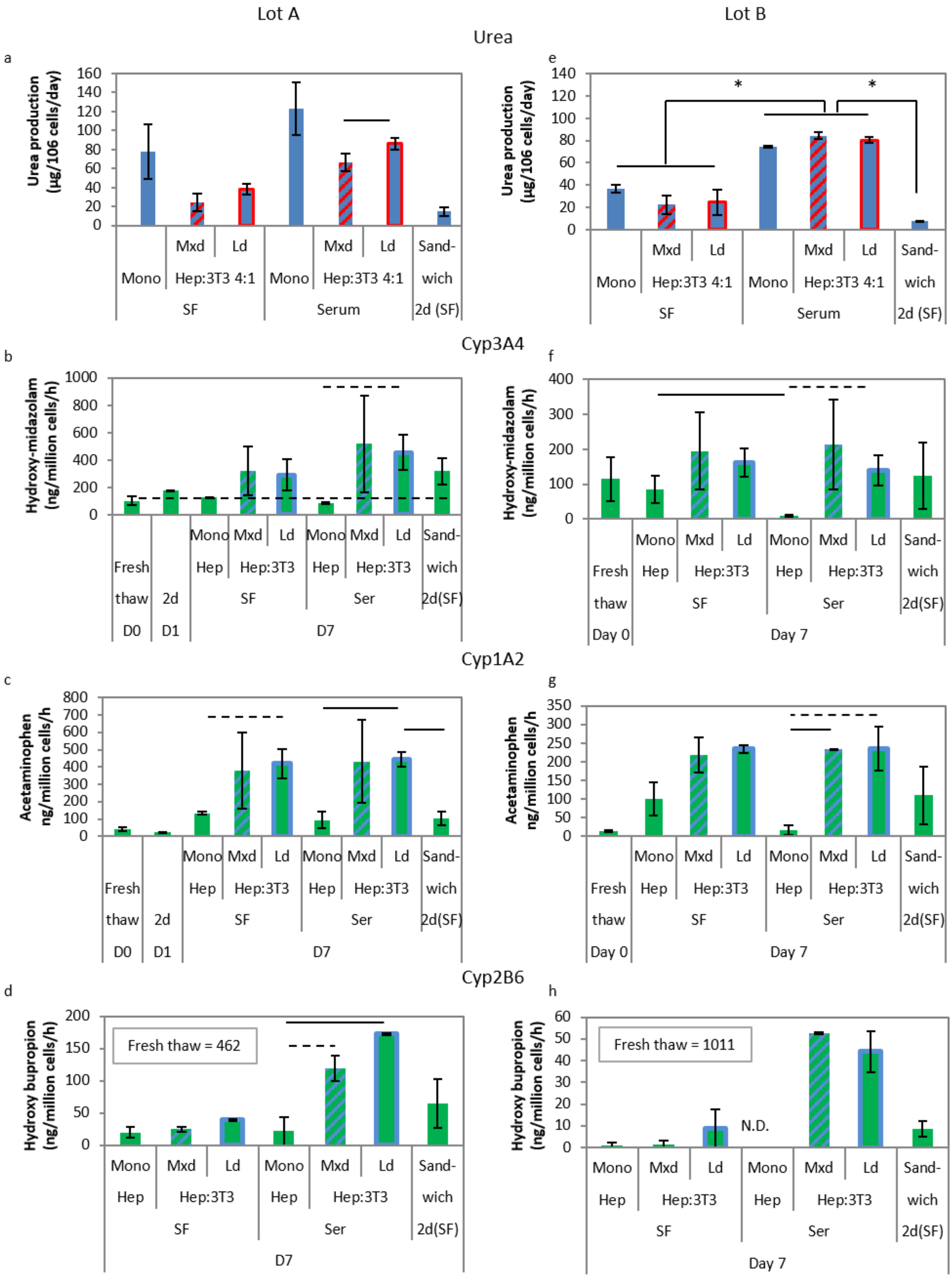


**Figure S7.** Function of mono- and co-culture spheroids for Lot A and B PHH on day 7. a and e. Urea production by mono- and co-culture spheroids on day 7 for Lot A (a) and Lot B (e). b, c, d. Cyp3A4, Cyp1A2, and Cyp2B6 activity for Lot A mono- and co-culture spheroids on day 7. Controls shown are the activity in freshly thawed cells (suspension) as well as hepatocytes in a collagen-matrigel sandwich. f, g, h. Cyp3A4, Cyp1A2, and Cyp2B6 activity for Lot B mono- and co-culture spheroids on day 7, along with suspension and sandwich values. Solid lines and asterisk indicate $p<0.05$. SF- serum-free medium, Ser – 10% serum-containing medium; Mxd – mixed co-culture; Ld – layered co-culture.
